# 2019-20 H1N1 clade A5a.1 viruses have better *in vitro* replication compared with the co-circulating A5a.2 clade

**DOI:** 10.1101/2023.02.26.530085

**Authors:** Nicholas J Swanson, Paula Marinho, Amanda Dziedzic, Anne Jedlicka, Hsuan Liu, Katherine Fenstermacher, Richard Rothman, Andrew Pekosz

**Author notes:** Corresponding author Andrew Pekosz, 615 North Wolfe Street, rm W2116, Baltimore, MD 21205.

## Abstract

Surveillance for emerging human influenza virus clades is important for identifying changes in viral fitness and assessing antigenic similarity to vaccine strains. While fitness and antigenic structure are both important aspects of virus success, they are distinct characteristics and do not always change in a complementary manner. The 2019-20 Northern Hemisphere influenza season saw the emergence of two H1N1 clades: A5a.1 and A5a.2. While several studies indicated that A5a.2 showed similar or even increased antigenic drift compared with A5a.1, the A5a.1 clade was still the predominant circulating clade that season. Clinical isolates of representative viruses from these clades were collected in Baltimore, Maryland during the 2019-20 season and multiple assays were performed to compare both antigenic drift and viral fitness between clades. Neutralization assays performed on serum from healthcare workers pre- and post-vaccination during the 2019-20 season show a comparable drop in neutralizing titers against both A5a.1 and A5a.2 viruses compared with the vaccine strain, indicating that A5a.1 did not have antigenic advantages over A5a.2 that would explain its predominance in this population. Plaque assays were performed to investigate fitness differences, and the A5a.2 virus produced significantly smaller plaques compared with viruses from A5a.1 or the parental A5a clade. To assess viral replication, low MOI growth curves were performed on both MDCK-SIAT and primary differentiated human nasal epithelial cell cultures. In both cell cultures, A5a.2 yielded significantly reduced viral titers at multiple timepoints post-infection compared with A5a.1 or A5a. Receptor binding was then investigated through glycan array experiments which showed a reduction in receptor binding diversity for A5a.2, with fewer glycans bound and a higher percentage of total binding attributable to the top three highest bound glycans. Together these data indicate that the A5a.2 clade had a reduction in viral fitness, including reductions in receptor binding, that may have contributed to the limited prevalence observed after emergence.

## Introduction

Seasonal influenza is a persistent contributor to global morbidity and mortality. While the burden of disease can vary significantly from year to year, overall influenza is responsible for millions of infections and hundreds of thousands of deaths annually^1,2^. There are several influenza virus strains and subtypes that vary in prevalence in different seasons. As these viruses circulate they accrue mutations, and some mutations contribute to antigenic drift and result in the need for regular updates to influenza vaccine formulations^3,4^. Mutations in the influenza virus can also result in changes to various aspects of viral fitness, including replication kinetics^5,6^, receptor binding^7-11^, and viral budding^12^. While antigenic structure and viral fitness are both important components of viral evolution, they are distinct phenomena; for example, a mutation mediating escape from preexisting immunity could simultaneously confer a detrimental effect on fitness. Studying how viruses balance these characteristics is crucial to improving our understanding of viral evolution.

The 2019-20 Northern Hemisphere influenza season was the third H1N1-predominant season since the 2009 H1N1pdm viruses emerged^13^. At the start of the season the 6B.1A.5a (A5a) clade constituted the majority of circulating H1N1, but it quickly dropped in prevalence as the A5a.1 and A5a.2 subclades emerged and began to cocirculate^14^ (Figure 1A). These clades differ by several amino acids on the viral hemagglutinin protein; A5a.1 contains D187A and Q189E while A5a.2 is defined by K130N, N156K, L161I, and V250A (Figure 1B). The clade-defining mutations for both subclades include mutations in canonical antigenic sites^15^, suggesting that they may have contributed to antigenic drift. Before its appearance in the 2019-20 season N156K had also emerged previously in regional circulation^16,17^, and had already been described as a potentially important mutation that changed antigenic structure in studies involving both ferret and human serum^16,18^. Both subclades also include mutations associated with the receptor binding site of hemagglutinin, including the 130 loop (K130N) and the 190 helix (D187A, Q189E). Changes to the receptor binding site can alter viral fitness by modifying the strength and diversity of receptor binding^7,10^. The N156K mutation, while not on the canonical receptor binding site, has also been associated with changes in receptor binding ^18^.

**Figure 1:**
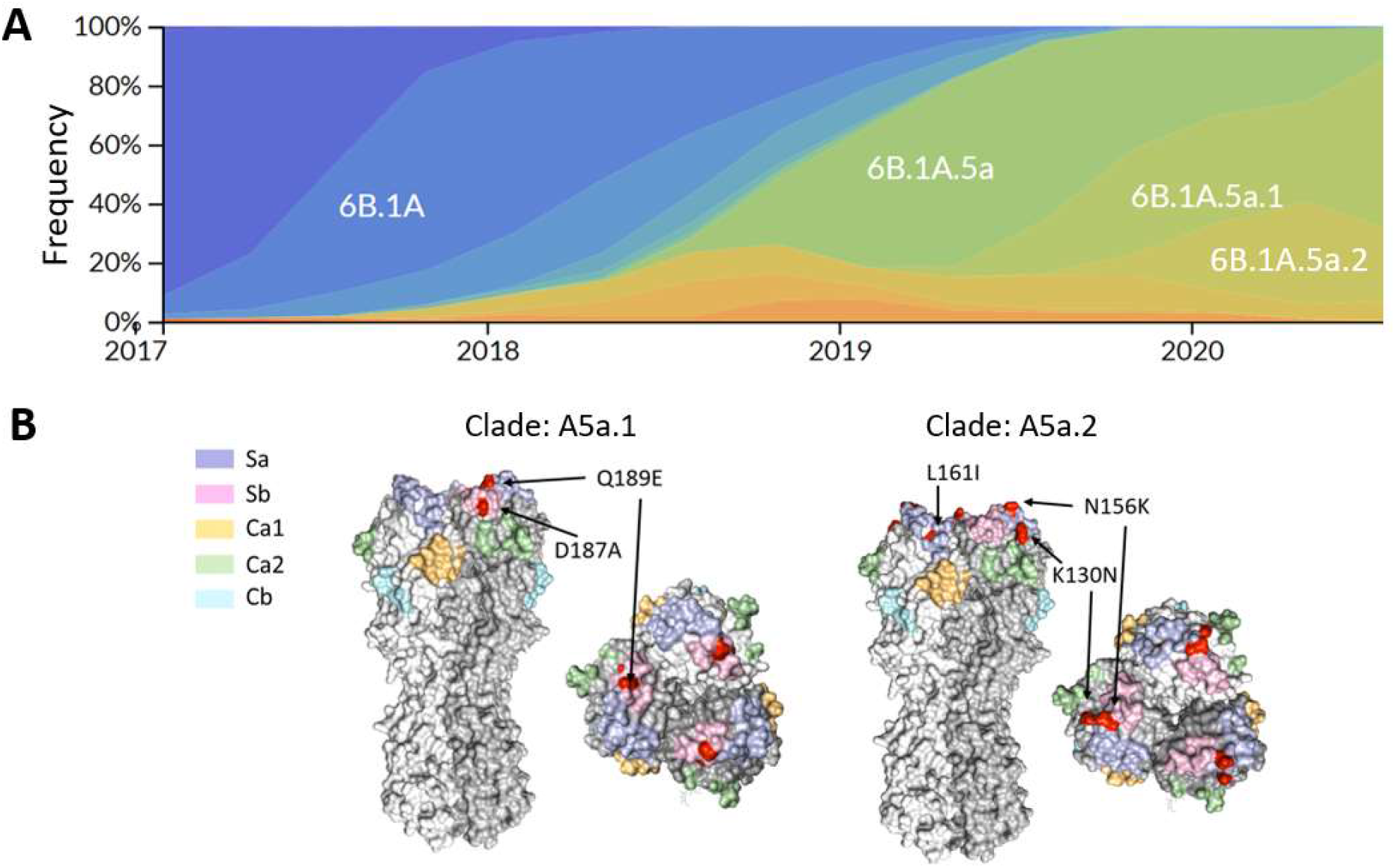
Emergence of the 6B.1A.5a.1 (A5a.1) and 6B.1A.5a.2 (A5a.2) clades of H1N1 and their clade-defining mutations. (A) Frequencies of H1N1 clades from January 2017-June 2020, generated through the NextStrain pipeline^1^. (B) Side and top view of hemagglutinin trimers from A5a.1 and A5a.2 clades are displayed with antigenic sites shaded using *PyMOL* (PDB: 3LZG). Clade-defining amino acid mutations are shown in red (not shown are V250A in HA1 and E179D in HA2 for the A5a.2 clade).

The H1N1 vaccine component for 2019-20 season was A/Brisbane/2/2018 which belongs to clade 6B.1A.1^14,19^, and a variety of studies have been performed to analyze the impact of the additional mutations on antigenic drift. Experiments using ferret serum post-infection with A5a or A5a.1 viruses showed a significant reduction in antibody recognition of the A5a.2 clade when compared with A5a or A5a.1, and ferret serum raised against A5a.2 viruses showed poor recognition of A5a.1 clade viruses^20-22^. Ferret antibodies have been noted to preferentially recognize antigenic site Sa and may therefore disproportionately respond to the A5a.2 mutations^23^; further antigenic characterization by the WHO using human serum post-vaccination with several prior vaccine strains showed a reduction in antibody recognition against both A5a.1 and A5a.2 viruses compared with A5a^20^. Vaccine effectiveness studies for the 2019-20 season, however, report a reduction in effectiveness against infection and hospitalization with A5a.2 viruses compared with A5a.1 and A5a^24-26^. Taken together, these data indicate that A5a.2 may have increased antigenic drift that allowed it to escape existing immunity.

Despite the potential antigenic advantages of the A5a.2 clade, A5a.1 predominated globally over both A5a.2 and A5a after its emergence and constituted over 90% of total circulating H1N1 by 2021^14^ (though it is noted that global circulation of influenza was greatly reduced during the 2020-2021 Northern Hemisphere influenza season due to the continuing SARS-CoV-2 pandemic). This suggests that factors of A5a.2 other than escape from preexisting immunity may have contributed to its lack of circulation. To investigate the phenotypic consequences of the clade-defining mutations of A5a.1 and A5a.2, representative viruses were isolated from influenza-positive patients in Baltimore, Maryland during the 2019-20 Northern Hemisphere influenza season. Growth curves, plaque assays, glycan arrays, and neutralization assays were performed, and A5a.2 was shown to have reduced replication, plaque formation, and receptor binding diversity compared with both A5a.1 and A5a. These data indicate that A5a.2 suffered a reduction in fitness that may have prevented it from becoming the predominant H1N1 clade in the 2019-20 Northern Hemisphere season.

## Results

A5a.1 and A5a.2 clade H1N1 viruses cocirculated in the 2019-20 influenza season in the Northern Hemisphere, with clade A5a.1 predominating (Figure 1A). A5a.1 and A5a.2 are distinguished by several distinct clade-defining mutations on the hemagglutinin protein (Figure 1B). A5a.1 contains D187A and Q189E mutations which occupy the 190 helix of the receptor binding site and antigenic site Sb. The A5a.2 clade is defined by K130N, N156K, L161I, and V250A. K130N is located on the 130-loop of the receptor binding site, while N156K and L161I occupy antigenic site Sa (Figure 1B). A5a.2 also contains E179D on the HA2 subunit of hemagglutinin. To characterize these clades, representative viruses were chosen from clades A5a.1, A5a.2, and the ancestral A5a clade. These viruses were collected from influenza-positive patients in Baltimore, Maryland during the 2018-19 and 2019-20 influenza seasons. For A5a.2 the virus A/Baltimore/R0675/2019 was chosen. For A5a.1 the viruses chosen were A/Baltimore/R0686/2019 and A/Baltimore/R0688/2019, with A/Baltimore/R0688/2019 containing the T72N mutation on the neuraminidase protein which resulted in an additional putative glycosylation site. A/Baltimore/R0496/2018 was chosen to represent the A5a clade. Table 1 depicts the complete set of amino acid differences between these viruses, with A/Baltimore/R0496/2018 as a reference.

**Table 1:** Amino acid differences between the 2019-20 viruses chosen for characterization compared with the 2018-19 A5a virus (A/Baltimore/R0496/2018). NP segment numbering was based on the established start codon for H3N2 and H1N1, instead of the recently emerged upstream start codon^2^.

|  | A/Baltimore/R0675/2019<br>(A5a.2) | A/Baltimore/R0686/2019<br>(A5a.1) | A/Baltimore/R0688/2019<br>(A5a.1) |
| --- | --- | --- | --- |
| HA1 | K130N/N156K/L161I/V250A | R45K/T120I/D187A/Q189E | D187A/Q189E |
| HA2 | E179D |  |  |
| NA | M19T/Y66F/N222K | S52N/I223V | I40T/S52N/T72N |
| M1 |  |  |  |
| M2 | H81Q | E70D/H81Q | E70D/H81Q |
| NP |  | A70S/V444I | A70S/V444I |
| NS1 | D189G | D189G | I145V/D189G |
| NS2 | I32V | I32V | I32V |
| PA | V63L/S225C/I505V | M336R/M374I | K213R/I465M/T528I |
| PA-X | V63L |  | S213G |
| PB1 | M174V/I728V | I728V | M646L/I728V |
| PB1-F2 |  |  |  |
| PB2 | Q183L/R630K | Q183L | Q183L |

Vaccinations in the 2019-20 Northern Hemisphere (NH) influenza season were shown to offer reduced protection against infection and hospitalization with A5a.2 compared with A5a.1^24-26^. Studies using ferret serum indicated that the N156K mutation resulted in antigenic drift and reduced recognition by antibodies raised by infection with the 2019-20 NH vaccine strain^20-22^, and experiments with human serum post-vaccination indicate a reduction in neutralizing titers against both A5a.1 and A5a.2^20^.

Experiments using human serum are an important component of antigenic characterization, as ferret antibodies have been shown to preferentially recognize antigenic site Sa compared with human antibodies^23^. To assess the antigenic consequences of clade-defining mutations on viral recognition by serum antibodies, neutralization assays were performed using serum from healthcare workers before and after receiving the 2019-20 NH influenza vaccine (Figure 2). Representative viruses were used from both the A5a.2 clade (A/Baltimore/R0675/2019) and the A5a.1 clade (A/Baltimore/R0688/2019). For pre- and post-vaccination timepoints, serum showed reduced neutralizing antibody titers against both circulating clades compared with the vaccine strain (A/Brisbane/02/2018) (Figure 2A). No significant differences in neutralizing titers were seen between A5a.1 and A5a.2 at either timepoint. Fold change in neutralizing antibodies were similar between all three viruses, with a comparable percent of participants seroconverting post-vaccination (Figure 2B). These data suggest that the clade-defining mutations of A5a.1 and A5a.2 showed similar escape from pre- and post-vaccination serum in this population that has a high annual acceptance rate of influenza vaccines.

**Figure 2:**
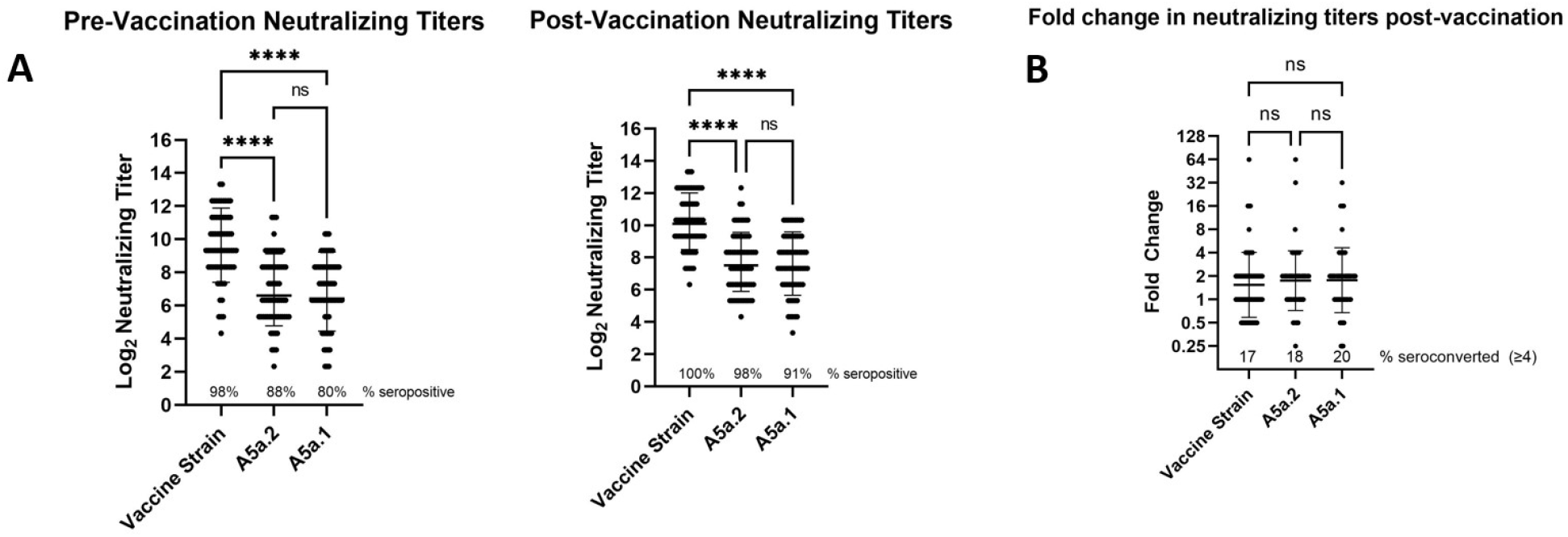
Comparison of neutralizing antibody titers in serum of healthcare workers pre- and post-vaccination. (A) Neutralizing titers of antibodies against the H1N1 vaccine strain (A/Brisbane/02/2018), A5a.2 (A/Baltimore/R0675/2019), and A5a.1 (A/Baltimore/R0688/2019). Neutralizing titers were significantly higher against the vaccine strain than the two circulating clades, and there were no significant differences between A5a.1 and A5a.2 pre- or post-vaccination. 1-way ANOVA with Tukey post-hoc, **** p<0.0001 (B) Fold-change of neutralizing titers post-vaccination show a similar seroconversion rate between all three viruses. Bars represent geometric mean, intervals are geometric standard deviation. One-way ANOVA with Tukey post-hoc. N=66

To assess differences in viral replication – which we will refer to as fitness - between these clades, growth curves and plaque assays were performed on the representative viruses from the A5a.1 and A5a.2 clades, as well as the ancestral A5a clade. Viral replication was assessed through low-multiplicity of infection (MOI) growth curves on MDCK-SIAT cells. These are MDCK cells stably transfected with human 2,6 sialtransferase, resulting in an overexpression of the canonical human influenza receptor (2,6 sialic acid) and a reduction in 2,3 sialic acid expression^27^. These growth curves revealed a reduction in A5a.2 viral titer at multiple timepoints compared with A5a and A5a.1 (Figure 3A). Growth curves were also performed on human nasal epithelial cell (hNEC) cultures, a primary cell culture model that is physiologically similar to the upper respiratory tract that seasonal influenza preferentially infect^5,28-33^. Growth curves on hNEC cultures reflected MDCK-SIAT growth curves, with A5a.2 producing significantly reduced titers at multiple timepoints including a sustained reduction in viral replication post-peak titer (Figure 3B). A5a and A5a.1 viruses both produced similar plaque sizes on Marin Darby Canine Kidney (MDCK) cells, while A5a.2 formed significantly smaller plaques (Figure 3C and 3D). Together these data indicate that the A5a.2 clade has reduced viral fitness compared with the cocirculating A5a.1 clade and its A5a precursor, shown through a reduction in plaque size and replication on multiple cell types.

**Figure 3:**
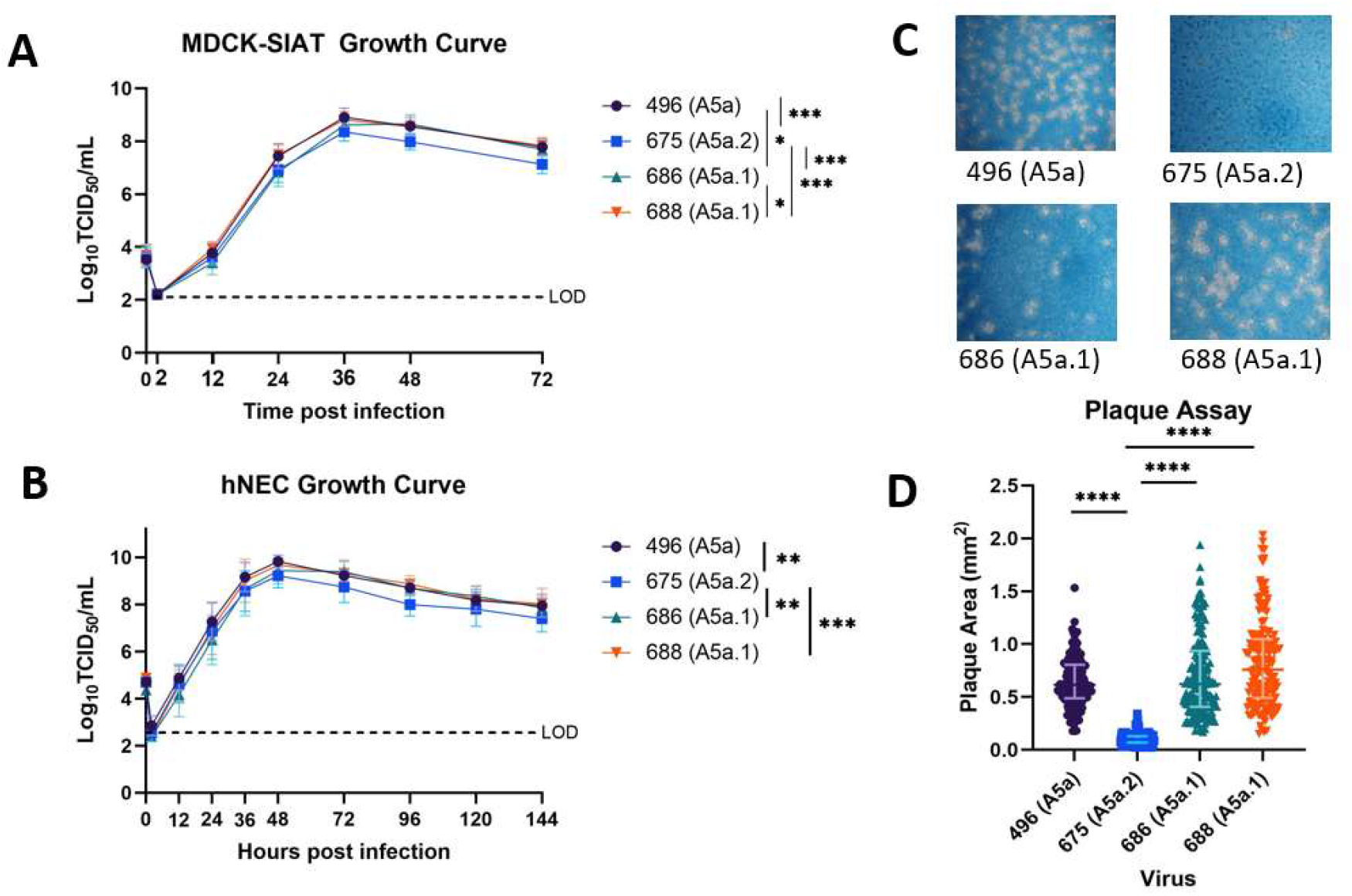
Characterization of Viral Fitness. (A) Growth curves on MDCK-SIAT cells (MOI 0.001) show a reduction in growth of A5a.2 viruses compared to A5a and A5a.1. (B) Growth curves on human nasal epithelial cells (MOI 0.01) show a similar phenotype, with A5a.2 having lower peak and post-peak titers. 2-way ANOVA with Tukey post hoc test, asterisks represent at least one time point with significant differences. * p<0.05, ** p<0.01, *** p<0.001. Each graph shows three experiments with four replicates each. (C) representative images of plaque sizes. D) Comparison of plaque areas reveal significantly reduced plaque sizes for the A5a.2 virus versus A5a and A5a.1 viruses. N=3, >50 plaques per virus. Kruskal-Wallis ANOVA with Dunn’s multiple comparisons test, **** p<0.0001.

Both A5a.1 and A5a.2 contain clade-defining mutations on the receptor binding site (RBS) of the hemagglutinin protein. Receptor binding is a crucial component of viral fitness and a reduction in receptor binding diversity or avidity could result in the reduced plaque sizes and replication seen for A5a.2. To assess whether these mutations resulted in phenotypic changes, receptor binding was assessed using the Consortium for Functional Glycomics glycan microarray (version 5.5) as previously described^7,34,35^. For all clades, viruses predominantly bound to glycans with alpha 2,6 linkages of the sialic acid with the penultimate sugar (Figure 4), which is the preferential linkage recognized by human influenza viruses^36,37^. To investigate differences in receptor binding diversity, percent of total binding was calculated for each glycan that bound with at least 5% of the relative fluorescence units (RFUs) of the highest bound glycan (Table 2).

**Table 2:**
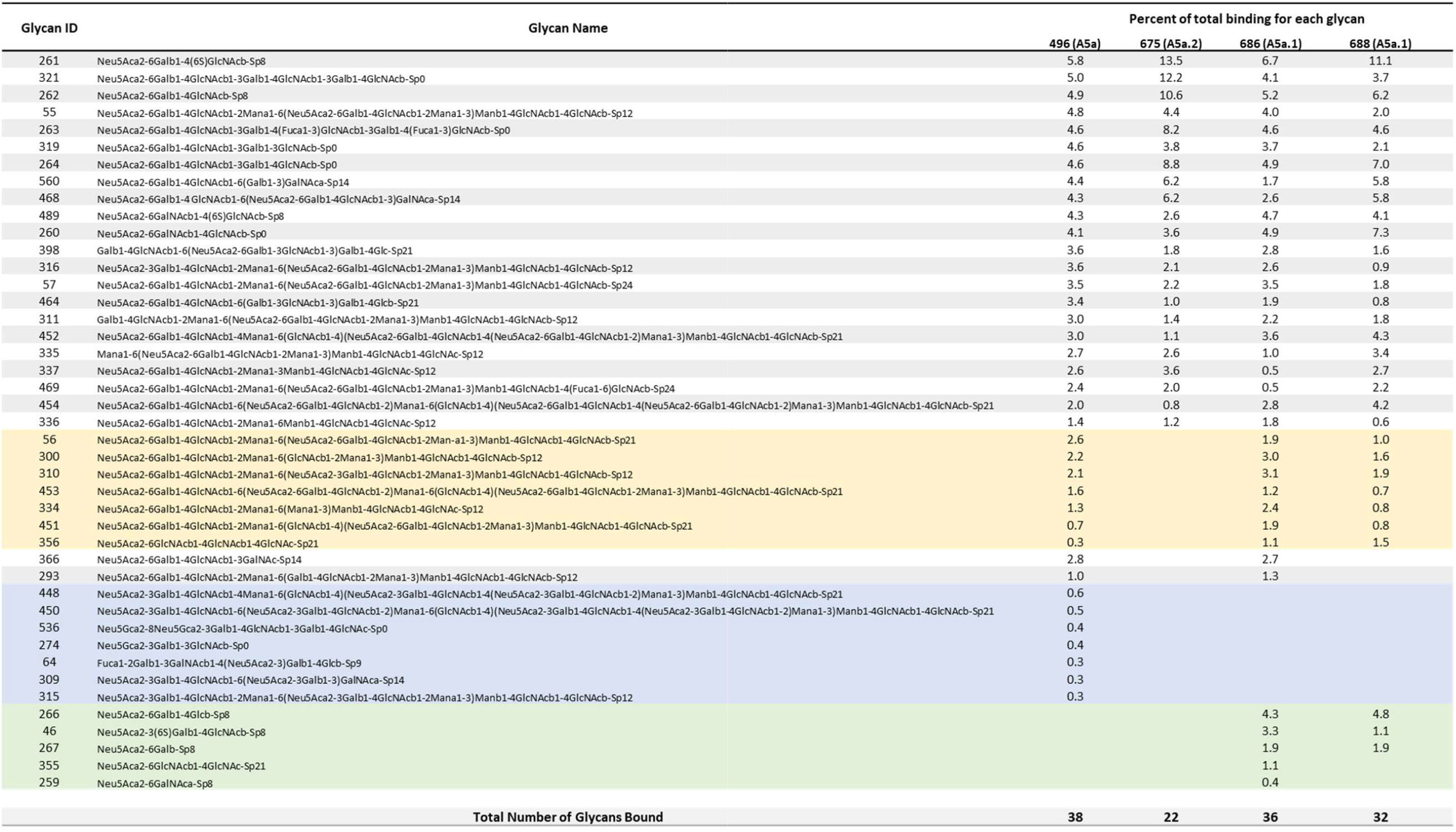
Percent of total glycan binding. Included in this list are all glycans with ≥ 5% of the RFU value of the highest bound glycan for that virus. For each virus, percent of total binding is listed for each glycan, and the total number of glycans bound for that virus is included at the bottom. Glycans highlighted in blue are bound only by the A5a clade virus, while glycans in green were bound only by viruses from the A5a.1 clade. Glycans highlighted in yellow were bound by every virus except for A5a.2.

**Figure 4:**
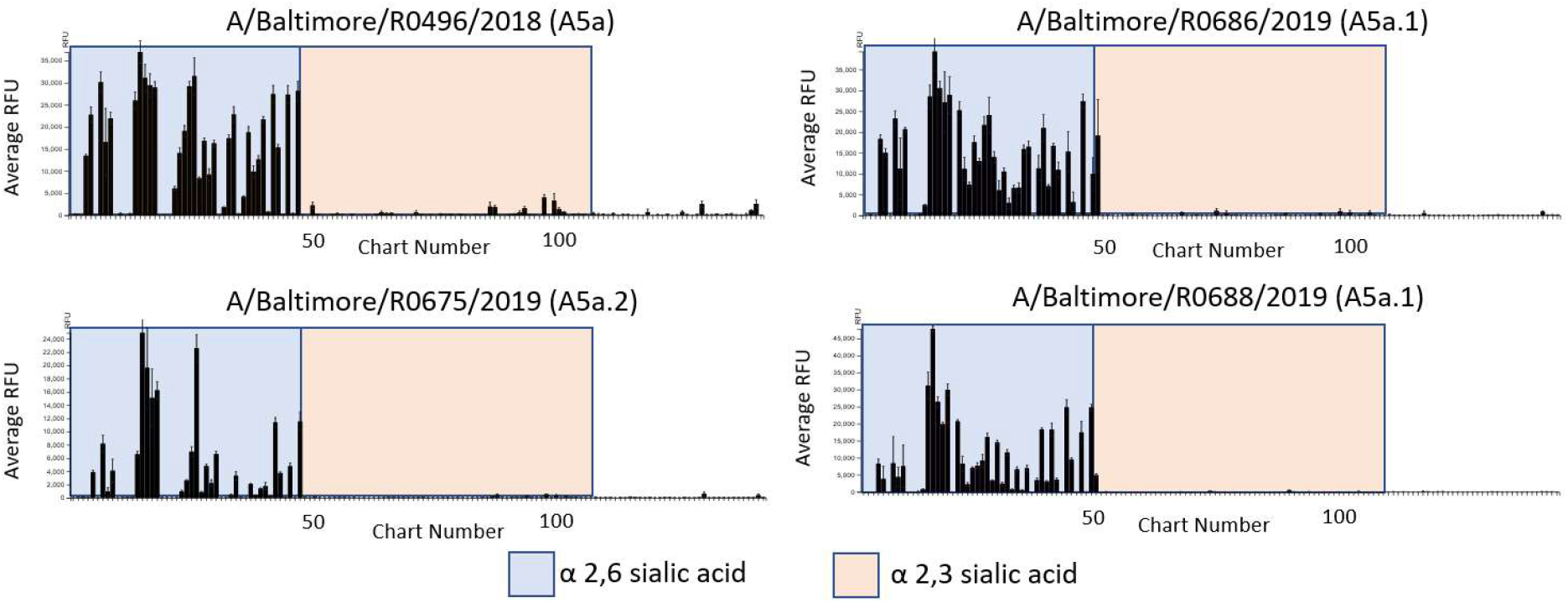
Consortium for Functional Glycomics (CFG version 5.5) glycan microarray profile of viral receptor binding of terminally sialylated glycans. Y axis depicts relative fluorescence units (RFUs), and X axis depicts individual glycans from the array.

The three highest bound glycans constituted over 35% of total binding for the A5a.2 virus, while the top three glycans account for 21% or less of total binding for A5a.1 and A5a viruses. Additionally, A5a and A5a.1 viruses bound to 32 or more glycans with at least 5% of the RFUs of the highest bound glycan, while A5a.2 only bound 22. Taken together these data indicate that viruses from the A5a.2 clade have reduced receptor binding diversity compared with the ancestral A5a and co-circulating A5a.1 clades. Both A5a (Figure 5) and A5a.1 (Figure 6) also demonstrated clade-specific binding to glycans not bound by the other clades; A5a bound to several branched alpha 2,3 SA glycans as well as Neu5Gc glycans not expressed by human cells (Figure 5), while A5a.1 bound to several single chain 2,6 SA glycans and one single chain 2,3 SA glycan (Figure 6). These data confirm that mutations in the receptor binding site can result in viral clades with distinct receptor binding profiles, and suggest that differences in receptor binding may have contributed to the fitness differences observed for A5a.2.

**Figure 5:**
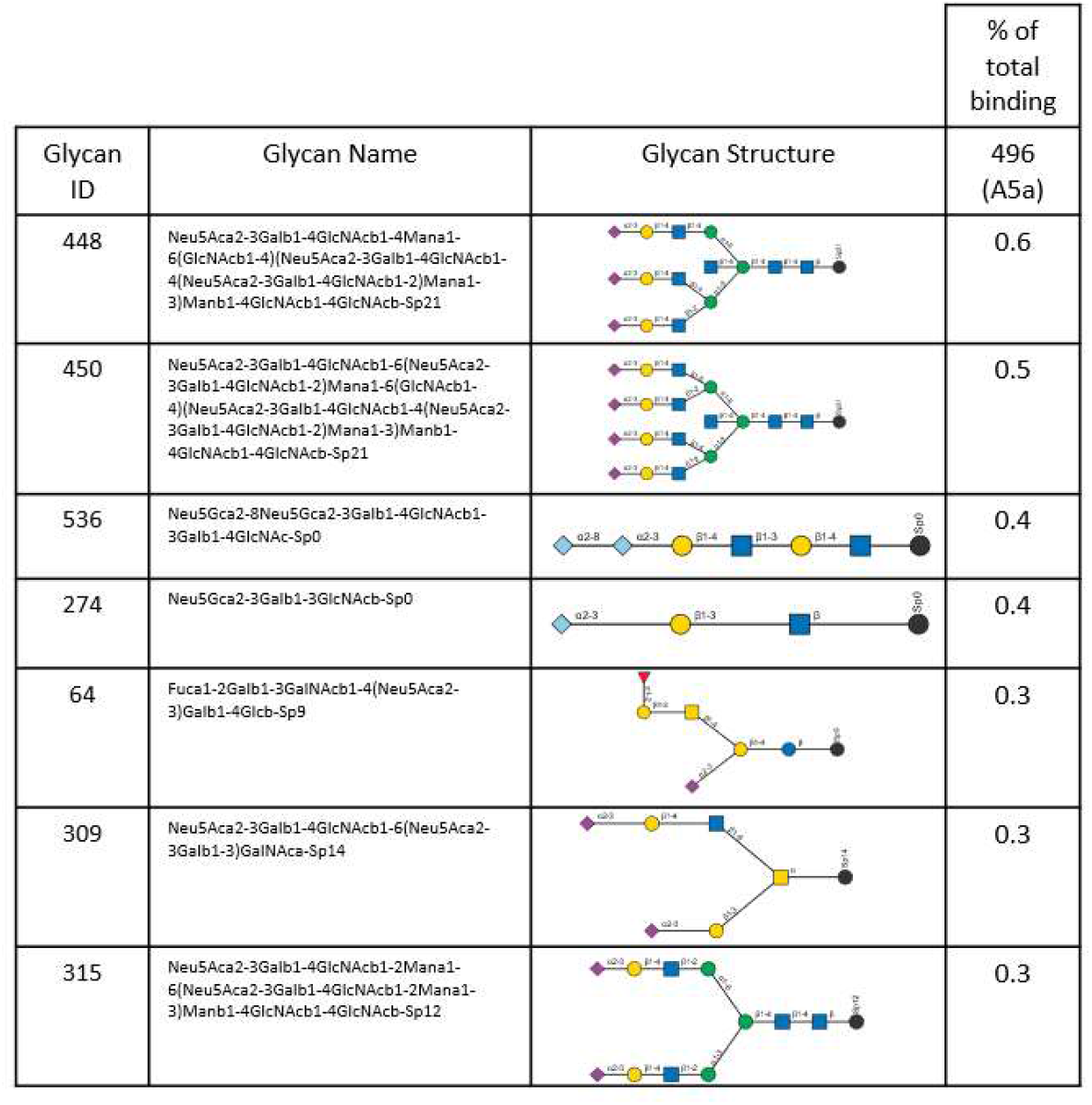
Glycans bound only by A5a virus.

**Figure 6:**
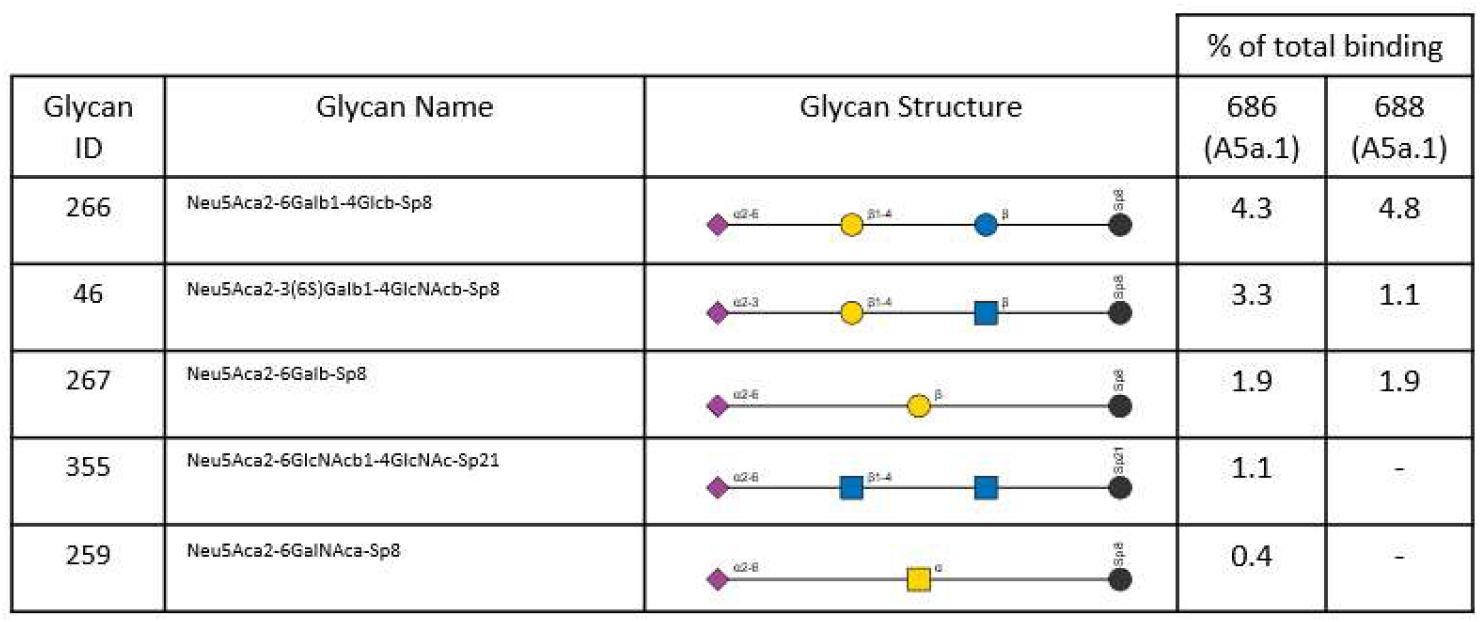
Glycans bound only by A5a.1 viruses.

## Discussion

The accumulation of mutations in circulating seasonal influenza results in antigenic drift and the need for annual vaccinations, as well as changes to viral fitness. Mutations in individual influenza proteins have been shown to affect plaque size^38^, viral replication^5,6,30,38^, and receptor binding preferences and diversity^7-11^. Importantly, there is overlap between functional and antigenic regions of both the neuraminidase (NA) and hemagglutinin (HA) proteins,^6,8,15,39^, and individual mutations have the potential to alter both antigenic structure and fitness simultaneously^6,40-43^. Antigenic differences in circulating clades have been associated with changes to viral receptor binding profiles^44^, and egg-adaptation mutations that changed receptor preferences in a live attenuated influenza virus also contributed to antigenic drift and reduced replication in human cells^38,45^.

The overlap between antigenic and functional regions may result in tradeoffs between novel mutations that improve one trait at the expense of the other. An example found in H3N2 is the emergence of a glycosylation site on the NA protein, which led to antigenic drift at the expense of viral replication and NA enzymatic activity^6,46^. This mutation increased in prevalence after its emergence, suggesting that the antigenic advantages may have offset any detriments to fitness^30^. Here we investigate a potential tradeoff between antigenic structure and fitness that led the A5a.1 clade of H1N1 to predominant in the 2019-20 Northern Hemisphere influenza season despite the apparent antigenic advantages of the cocirculating A5a.2 clade. We show that A5a.2 had deficits in plaque size, replication, and receptor binding compared to A5a and A5a.1 that may have contributed to its limited prevalence in the 2019-20 season.

A5a.1 and A5a.2 are distinguished by several important mutations to the viral hemagglutinin segment. A5a.1 is defined by D187A and Q189E, while A5a.2 viruses have the mutations K130N, N156K, L161I, and V250A as well as the HA2 subunit mutation E179D. The N156K mutation of A5a.2 has emerged independently in earlier seasons and has been shown to cause antigenic drift in studies using both ferret and human serum^16,18^, and previous antigenic characterization of the 2019-20 NH clades has shown that A5a.2 is drifted from both A5a.1 and the 2019-20 NH vaccine strain in ferret serum and vaccine effectiveness studies^20-22,24-26^. Experiments with human serum show a similar degree of antigenic drift in both A5a.1 and A5a.2 compared with preceding clades^20^.

To assess antigenic differences between the 2019-20 NH H1N1 clades, neutralization assays were performed on representative A5a.1 and A5a.2 viruses using serum from healthcare workers from the 2019-20 NH influenza season. These experiments revealed a similar reduction in neutralization of both A5a.1 and A5a.2 viruses compared with the vaccine strain (Figure 2), both pre- and post-vaccination. Fold-increase in neutralizing titers after vaccination were similar for all three viruses. It is important to note that healthcare workers represent a highly vaccinated population, with mandatory vaccination leading to over 90% vaccine coverage^47^. Consecutive yearly vaccination has been shown to increase pre-vaccination antibody titers^48^, and likely contributed to the high pre-vaccination neutralizing antibody levels observed here. High pre-vaccination antibody titers have previously been inversely correlated with seroconversion after vaccination^49,50^, and it is possible that the increased antigenic drift of A5a.2 reported elsewhere may have been masked in our study by high baseline antibody levels.

While antigenic comparisons of these clades have revealed similar or increased antigenic drift in A5a.2, the predominance of A5a.1 in the 2019-20 NH influenza season suggests a potential fitness advantage over A5a.2. Here we show that a representative A5a.2 virus produces significantly smaller plaques in MDCK cells compared with viruses from both A5a.1 and the ancestral A5a clade. Small viral plaque size can be caused by multiple factors^51^, and may be influenced by many aspects of the viral life cycle including replication, receptor binding, or cell-to-cell spread. To assess differences in viral replication, growth curves were performed in MDCK-SIAT and human nasal epithelial cell (hNEC) cultures. MDCK-SIAT cells are an immortalized cell line that overexpress alpha 2,6 sialic acids, the canonical human influenza receptor^27^, and hNEC cultures are a primary cell culture system that models the upper respiratory tract^52^. hNEC cultures are grown at an air-liquid interface and differentiate into multiple epithelial cell types including ciliated and mucus-producing cells^28,52^. While MDCK-SIAT cells are engineered to be easily infected with human influenza, hNEC cultures are considered to be more physiologically relevant and may reveal differences not seen in MDCK-SIAT growth curves^6,29,30^. Both growth curves were performed at 33°C to reflect the temperature of the upper respiratory tract. In both MDCK-SIAT and hNEC growth curves A5a.2 consistently demonstrated a significant reduction in replication compared to A5a.1 or A5a (Figure 3). This reduced replication could have contributed to the small plaque sizes of A5a.2.

To further explore the potential mechanisms of reduced fitness in A5a.2, glycan arrays were performed on the representative viruses from the three clades. These experiments measure viral binding to a variety of polysaccharides, and can reveal differences in both receptor preferences and binding diversity. They can also provide insight into host specificity; the canonical influenza receptors are glycans with a sialic acid (SA) as their terminal sugar, and different SA linkages to the penultimate sugar are differentially preferred by avian and human influenza viruses. Human influenza preferentially binds to alpha 2,6 SA linkages while avian influenza recognizes alpha 2,3 SA^36,37^. Glycan arrays on the representative viruses confirm a preference for alpha 2,6 SA glycans for all viruses tested, but there was considerable difference in binding diversity between clades. A5a.1 and A5a viruses bound to over 45% more glycans than A5a.2 with at least 5% of the relative fluorescence units of the highest bound glycan (Table 2). Additionally, the top three bound glycans represent over 35% of total binding for A5a.2, while the top three only constituted 21% or less of total binding for A5a.1 and A5a viruses.

Differences in receptor binding have previously been associated with reduced replication kinetics^7^ and virus propagation^10^, though this correlation does not appear in every context^35,53-55^. Diversity of receptor binding and low-affinity receptor interactions have also been shown to be important for viral entry into cells^56^, and it is possible that the reduction in binding diversity seen in the A5a.2 clade could impact its fitness through impaired cell binding and entry. Additionally, viruses from both A5a and A5a.1 clades each bound to unique glycans not bound by the other clades (Figures 5 and 6). This confirms that different clades can have unique binding profiles that distinguish them from each other, and further demonstrates the reduced binding diversity of A5a.2 which had no uniquely bound glycans.

Interestingly, as A5a.2 has continued to circulate it quickly gained a series of mutations on the hemagglutinin protein, including K54Q, A186T, Q189E, E224A, R259K, and K308R, and has subsequently become the predominant circulating clade of H1N1^14^. Several of these mutations occupy both antigenic and receptor binding sites, suggesting that they may have served some compensatory function that allowed the virus to improve its fitness and/or antigenic structure and outcompete A5a.1. Further investigation will be needed to identify the impact of these more recent mutations on the viral phenotype. It is also important to note that this study characterized clinical isolates of viruses representing the various clades, and differences between these viruses are not limited to the hemagglutinin (HA) protein (Table 1). Additional research is needed to identify the role of individual clade-defining HA mutations on the overall reduction in fitness of A5a.2. These findings nevertheless highlight the deficiencies in receptor binding and viral fitness that may have contributed to the limited spread of the A5a.2 clade in the 2019-20 Northern Hemisphere influenza season.

## Methods

### Ethics statement and human subjects

Serologic samples for this study were obtained from healthcare workers (HCWs) recruited from the Johns Hopkins Centers for Influenza Research and Surveillance (JHCEIRS) during the annual Johns Hopkins Hospital (JHH) employee influenza vaccination campaign in the Fall of 2019. Pre- and post-vaccination (∼28 day) human serum were collected from subjects, who provided written informed consent prior to participation. The JHU School of Medicine Institutional Review Board approved this study, IRB00288258. Virus was isolated for this study from deidentified influenza A virus H1N1 positive samples, collected from patients who provided written informed consent during the 2019-20 influenza season at the Johns Hopkins Hospital, under the JHU School of Medicine Institutional Review Board approved protocol, IRB00091667.

### Cell Cultures

Madin-Darby canine kidney (MDCK) cells and MDCK-SIAT cells were maintained in complete medium (CM) consisting of Dulbecco’s Modified Eagle Medium (DMEM) supplemented with 10% fetal bovine serum, 100 units/ml penicillin/streptomycin (Life Technologies) and 2mM Glutamax (Gibco) at 37°C and 5% CO_2_. Human nasal epithelial cells (PromoCell) were seeded on 24-well Corning transwell plates with PneumaCult Ex-Plus media on apical and basolateral sides. After cells reached confluence (approximately 10 days, determined by a trans-epithelial electrical resistance reading of >300 ohms), media was switched to PneumaCult ALI media and the apical surface was left at an air-liquid interface (ALI). hNEC cultures were considered fully differentiated when mucus and beating cilia were visible, approximately three weeks after transition to ALI.

### Viral Isolation and Sequencing

Nasopharyngeal swabs or nasal wash from individuals who were influenza A positive during the 2019-20 Northern Hemisphere influenza season were used for virus isolation on primary cells. The apical side of hNEC wells were washed twice with 300ul of phosphate buffered saline (PBS) and 100ul of sample was added to the cells and incubated for two hours. The sample was then aspirated and cells were washed twice with 300ul of PBS. At three, five, and seven days post-infection 300ul of hNEC infection media (DMEM supplemented with .3% BSA (Sigma), 100 units/ml pen/strep (Life Technologies), 2mM Glutamax) was added to the well and incubated for ten minutes. TCID50 was performed on collected media and stocks were made from the collected media when virus was detected at concentrations greater than 10^4 TCID50/mL.

To generate viral stocks, T75 flasks (Corning) were seeded with MDCK-SIAT cells and grown to confluence in complete media. Viral isolates from the previous step were diluted to a multiplicity of infection (MOI) of 0.001 in infection media (hNEC infection media plus 5μg/ml N-acetyl trypsin). Cells were washed twice with PBS plus 100mg/L each of anhydrous calcium chloride and magnesium chloride hexahydrate (PBS +/+) and inoculum was added to the flasks and incubated at 33°C, rocking every 15 minutes. Inoculum was then aspirated, and 13mL of infection media was added to the flask. After flasks showed 75% cell death (∼3 days post infection), media was collected and centrifuged at 500g for 10 minutes to remove cell debris. Stocks were aliquoted and stored at -80 °C.

Viral RNA was extracted using the QIAamp viral RNA mini extraction kit, and Illumina RNA Prep with Enrichment(L) Tagmentation was used for library preparation. Quality was checked using the Qubit and Agilent Bioanalyzer, and samples were sequenced with using a MiSeq Illumina sequencer. Consensus sequences were generated using the DRAGEN RNA Pathogen Detection pipeline.

Tissue Culture Infections Dose 50 (TCID50)

96-well plates were seeded with MDCK-SIAT cells. After reaching confluence, plates were washed twice with 100ul per well of PBS +/+ and 180ul of infection media was added to each well. Ten-fold serial dilutions were performed on viruses and 20ul of the dilution was added to its corresponding well in the cell plate. Each virus dilution was added to cell plates in sextuplicate, plates were incubated for six days at 33°C before being fixed with 4% paraformaldehyde for >3 hours and stained overnight with naphthol blue-black. 50% tissue culture infectious dose was calculated as previously described^57,58^.

### Plaque Assay

MDCK cells were grown in complete medium to confluence in 6-well plates. After reaching confluence, media was removed and cells were washed twice with PBS +/+. Ten-fold serial dilutions of viruses were made in infection media, and 250uL of the virus dilutions were added to the wells. Plates were incubated for one hour at 33°C and rocked every 15 minutes to ensure even distribution of inoculum. After one hour, the virus inoculum was removed and phenol-red free DMEM supplemented with .3% BSA (Sigma), 100U/ml pen/strep (Life Technologies), 2mM Glutamax (Gibco), 5mM HEPES buffer (Gibco) 5μg/ml N-acetyl trypsin (Sigma) and 1% agarose was added. Cells were incubated at 33°C for two days and then fixed with 4% paraformaldehyde overnight. After fixing, the agarose overlay was removed and cells were stained with naphthol-blue black. Plaque area was analyzed in ImageJ.

### Viral Growth Curves

For MDCK-SIAT growth curves, cells were seeded in a 24-well plate. After cell plates reached confluence, they were washed twice with PBS +/+ (described above). Virus inoculum was made by diluting virus stock to an MOI of 0.001 in infection media (described above). Inoculum was added to the cell wells and incubated at 33°C for one hour with plates being rocked every 15 minutes. After one hour, inoculum was removed and plates were washed twice with PBS +/+ and 500ul of infection media was added to each well. For each timepoint, infection media was collected from the wells and replaced with 500ul of fresh infection media. Viral titer in collected media was determined by TCID50 (described above).

hNEC growth curves were performed on fully differentiated hNECs grown at an air-liquid interface. Virus inoculum was made by diluting virus stocks to an MOI of 0.01 in hNEC infection media. Wells were washed three times with hNEC infection media and inoculum was added to the wells. Inoculum was incubated on the cells for two hours, after which it was removed and cells were washed three times with PBS and left at an air-liquid interface. For each timepoint, 100ul of hNEC infection media was added to the wells and incubated at 33°C for 10 minutes. Media was then collected and cells were left at ALI. Virus concentration in collected media was determined through TCID50.

### Partial virus Purification and Labeling with Alexa Fluor 488

T150 flasks were seeded with MDCK-SIAT cells and grown to confluence in complete media. Virus stocks were diluted to 0.001 MOI in infection media. Cells were washed twice with PBS +/+ and inoculum was added to the flasks and incubated at 33°C, rocking every 15 minutes. Inoculum was then aspirated, and 18mL of infection media was added to the flasks. After cells showed 75% death (∼3 days post infection), media was collected and centrifuged at 500g for 10 minutes to remove cell debris.

5mL of 20% sucrose in PBS was added to Ultra-Clear ultracentrifuge tubes (Beckman Coulter), and the remaining volume of the tube was filled with the collected infection media from the previous step.

Samples were centrifuged at 24,800 RPM for 1 hour, after which the liquid was aspirated and the pellet was resuspended in 100ul of PBS +/+. TCID50 was performed on purified viruses to confirm activity and concentration.

1mg of Alexa Fluor 488 (Invitrogen A20000) was resuspended in 1mL of diH_2_O. 25ul of Alexa Fluor solution was added to 200ul of purified virus and 20ul of 1M NaHCO3 (pH 9.0), mixed by pipetting, and incubated in the dark at room temperature (23 °C) for 1 hour. Solution was added to Slide-A-Lyzer™ 7K MWCO MINI Dialysis Devices (Thermo 69560), devices were loaded onto dialysis floats (Thermo 69588) and dialyzed while stirring at 4 °C in 1L PBS. After 1 hour, PBS was replaced with 1L of fresh pre-chilled PBS and the dialysis was continued overnight. The next morning, the PBS was again replaced and the dialysis was continued for another hour. Dialyzed labeled virus was collected and stored at -80 °C, and TCID50 was performed to confirm virus activity and concentration. Titer of labeled viruses were: 2.32E+10 TCID50/mL (A/Baltimore/R0496/2018), 5.00E+09 TCID50/mL (A/Baltimore/R0675/2019), 2.32E+10 (A/Baltimore/R0686/2019), 7.34E+10 TCID50/mL (A/Baltimore/R0688/2019).

### Glycan Array

The microarray chosen was version 5.5 of the CFG glycan microarray, which includes 562 glycans in replicates of 6. Glycan microarray slides were submerged in 100 milliliters of PBS wash buffer (phosphate-buffered saline containing 0.005% Tween-20) for five minutes to re-hydrate. Samples were diluted 1:100 and in PBS binding buffer (PBS with 0.05% Tween-20 and 1% bovine serum albumin), and 70 microliters of diluted sample were loaded onto the slide. A cover slip was placed over the slide and incubated at room temperature for one hour in darkness, after which the cover slip was removed and the slide was washed four times in 100 milliliters of PBS wash buffer, then PBS, then deionized water. Slides were then dried before being read in a GenePix microarray scanner.

For each set of six replicates, only the median four values were used in analysis. A graph of the raw data was visually analyzed to confirm selective binding of terminally sialylated glycans, after which data was filtered to only include these glycans and organized by structure using the Glycan Array Dashboard (GLAD)^59^. Glycans were considered in final analysis if virus bound to them with at least 5% of the relative fluorescence units (RFUs) of the highest bound glycan; total binding was found by summing the RFUs of all of these glycans, and percent of total binding was found by dividing the RFUs of individual glycans by total binding. Structures of glycans uniquely bound by one clade were identified using GLAD.

### Serum Neutralization Assay

Pre- and post-vaccination human serum obtained through the Johns Hopkins Center for Excellence in Influenza Research and Surveillance (JH-CEIRS) study (HHSN272201400007C) were used in this study. Lyophilized Receptor Destroying Enzyme II (RDE, Hardy Diagnostics) was dissolved into 20 mL of saline (0.9% NaCl in H_2_O) and a 1:3 ratio of serum and RDE was incubated at 37°C overnight followed by RDE inactivation at 57°C for 35 minutes. MDCK cells were seeded on 96-well plates. When cells reached confluence plates were washed twice with 100ul per well of PBS +/+. Virus inoculum was prepared by diluting virus stock in infection media to a concentration of 1,000 TCID50/ml. RDE-treated serum was serially diluted twofold in 96-well U bottom plates (Thermo) using infection media yielding 55ul per dilution, and 55ul of virus inoculum was added to each dilution. The serum/virus mixture was incubated at room temperature for one hour, and then 50ul of serum/virus mixture was transferred in duplicate onto the cell plates, yielding 25 TCID50 per well. After 24 hours, the serum/virus mixture was removed and plates were washed once with PBS +/+ before adding 100ul per well of infection media. Six days post-infection the plates were fixed with 4% paraformaldehyde for >3 hours and stained overnight with naphthol blue-black.

### Statistical Analyses

All statistical analyses were performed in GraphPad Prism 9.1.0. Growth curves were analyzed using 2-way ANOVA with Tukey post-hoc test. Plaque assays were analyzed using Kruskal-Wallis ANOVA with Dunn’s multiple comparison test. Serology was analyzed using either 2-way ANOVA or 1-way ANOVA, both with Tukey post-hoc test.

### Data availability

All data is available upon request.

## Acknowledgements

This work was supported by N272201400007C and N7503021C00045 for the Johns Hopkins Centers of Excellence in Influenza Research and Research as well as the Richard Eliasberg Family Foundation. We acknowledge the Protein-Glycan Interaction Resource of the CFG and the National Center for Functional Glycomics (NCFG) at Beth Israel Deaconess Medical Center, Harvard Medical School (supporting grant R24GM137763) for glycan microarray studies. The authors thank the healthcare workers who enrolled and participated in the Johns Hopkins Center for Excellence in Influenza Research and Surveillance study. We are grateful for the efforts of the clinical coordination team at JHH who collected samples. We thank the laboratories of Sabra Klein, Kimberly Davis, Nicole Baumgarth, and Andrew Pekosz for discussion of data and future directions.

